# Modular development enables rapid design of media for alternative hosts

**DOI:** 10.1101/2021.04.30.442183

**Authors:** Andrew M. Biedermann, Isabella R. Gengaro, Sergio A. Rodriguez-Aponte, Kerry R. Love, J. Christopher Love

## Abstract

Developing media to sustain cell growth and production is an essential and ongoing activity in bioprocess development. Modifications to media can often address host or product-specific challenges, such as low productivity or poor product quality. For other applications, systematic design of new media can facilitate the adoption of new industrially relevant alternative hosts. Despite manifold existing methods, common approaches for optimization often remain time and labor intensive. We present here a novel approach to conventional media blending that leverages stable, simple, concentrated stock solutions to enable rapid improvement of measurable phenotypes of interest. We applied this modular methodology to generate high-performing media for two phenotypes of interest: biomass accumulation and heterologous protein production, using high-throughput, milliliter-scale batch fermentations of *Pichia pastoris* as a model system. In addition to these examples, we also created a flexible open-source package for modular blending automation on a low-cost liquid handling system to facilitate wide use of this method. Our modular blending method enables rapid, flexible media development, requiring minimal labor investment and prior knowledge of the host organism, and should enable developing improved media for other hosts and phenotypes of interest.

## Introduction

Achieving high volumetric productivities of biologic drugs in cultivation is a key step in advancing candidate biologic drugs. The outcome of this effort ultimately impacts manufacturing costs as well as readiness for transitioning clinical-stage development (Love, Love, & Barone, 2012). The development of standard, chemically defined media for established manufacturing hosts, such as CHO, has made such transitions efficient for monoclonal antibodies by achieving high biomass accumulation, cell viability, operational consistency, and specific productivities, streamlining development efforts (McGillicuddy, Floris, Albrecht, & Bones, 2018; Rodrigues, Costa, Henriques, Azeredo, & Oliveira, 2012). Nonetheless, optimizing productivity or quality attributes for a specific product often still requires further refinement of media (Ritacco, Wu, & Khetan, 2018). Such development may require evaluating dozens of variants derived from a common standard formulation to address the specific challenges encountered (Gagnon et al., 2011; Loebrich et al., 2019). Media development for entirely new biomanufacturing technologies, such as alternative hosts (Matthews, Kuo, Love, & Love, 2017a) or new product modalities (Lu et al., 2016), may also require new formulations or extensive optimizations due to limited prior knowledge.

Common approaches to develop a medium to optimize a phenotype of interest are often labor intensive, low throughput, or rely heavily on extensive analytical capacity (Galbraith, Bhatia, Liu, & Yoon, 2018). For example, analysis of residual media after cultivation requires extensive capabilities for analytical characterization and prior experience with the manufacturing host to identify potentially limiting or toxic media components (Mohmad-Saberi et al., 2013; Pereira, Kildegaard, & Andersen, 2018). As a result, optimizations can be slow and iterative. Furthermore, for an alternative host such as *Komagataella phaffii* (formerly known as *Pichia pastoris*), there is substantially less, if any, prior knowledge available to establish profiles for residual components in media after fermentation. Other analytical techniques like RNA-seq combined with methods for reporter metabolite analysis can guide media optimization, to generate testable hypotheses regarding beneficial modifications to media (Matthews et al., 2017a). Such genome-scale approaches, however, require prior host-specific knowledge, such as well-annotated genomes, and are still limited by slow iteration and labor-intensive preparations of new media to test the hypotheses generated from computational analyses.

Alternative strategies for blending basal components for media allow linear combinations of existing media to explore many variations rapidly (Jordan, Voisard, Berthoud, & Tercier, 2013). While this approach avoids slow iterative analyses, the typical experiment is labor intensive to perform, often requiring independent preparations of over a dozen stock media to combine (Rouiller et al., 2013). Similar to analytical-based approaches for optimization, the selected variations of media are simultaneously guided and constrained by prior experience and media designs, which may limit the breadth of components examined (Kennedy & Krouse, 1999). For less established hosts with fewer available formulations of media, media blending may also require fully *de novo* formulations for initial studies. Further complicating such designs, different and new components for media can present challenges in solubility or unanticipated interactions with other elements in the formulations (Ritacco et al., 2018). New approaches to blending could, however, enable fast, flexible experimentation and minimize the time, labor, and analytical development needed initially to optimize media for new applications and phenotypes.

Here, we present a novel and generalizable approach for the modular development of media and demonstrate its use to create optimized media for two different phenotypes—cellular growth and recombinant expression of a protein (as measured by the secreted heterologous protein titer) from *Pichia pastoris*. Our approach comprises two modular parts for blending and optimization. We determined that a set of simple concentrated stock solutions constructed in defined modules could generate many media by blending or dilution. We then automated a simple, inexpensive liquid handling system (Opentrons OT-2) to enable high-throughput screening for the effects of diverse media on a phenotype of interest in milliliter-scale batch cultures. To maximize the benefit of this automated blending, we also developed an algorithmic framework for systematic modular media optimization, beginning from a simple minimal media (here a YNB-based one). This framework provides insights pertaining to key media components during stages of optimization, as well as overall mapping of the design space for the media. In the examples presented here, the resulting defined media developed with this strategy outperformed commonly used BMGY and BMMY complex formulations for biomass accumulation and secreted heterologous protein production.

## Materials and Methods

### Strains and cultivation conditions

Media for evaluating biomass accumulation were developed using a previously described strain expressing G-CSF under control of the pAOX1 promoter (Crowell et al., 2020). 24-well plate screens were conducted as described previously, except cells were grown on a labForce shaker and were only sampled 24 hours after inoculation (Matthews et al., 2017a). BMGY, BMMY, and RDM media were formulated as described previously for shake flasks (Matthews et al., 2017a). All cultivations were inoculated from a working cell bank at an initial cell density of 0.1 OD/mL. For each working cell bank, cells grown in 1 L shake flasks with a 200 mL working volume of RDM were harvested during exponential growth (4-5.5 OD/mL) via centrifugation at 1500 rcf for 4 minutes at 23°C and resuspended in an equal volume mixture of RDM and 50 v/v% glycerol. This mixture was then distributed into 700 μL aliquots and stored at −80°C, resulting in a cell density of ~30 OD/mL for the cell bank.

Media for evaluating enhanced production were developed using a strain expressing a rotavirus-derived subunit vaccine candidate, P[8], under the control of the pAOX1 promoter described previously (Dalvie et al., 2020). Biomass accumulation proceeded for 24 hours; cells reached an initial induction density of ~20 OD/mL. Cultures were then exchanged into production media and allowed to produce protein for an additional 24 hours. Supernatant was harvested by centrifugation at 1500 rcf for 4 minutes at 23°C and filtered using a Captiva 96 well 0.2 μm filter plate (Agilent Technologies, Santa Clara, CA) prior to titer measurement by RP-UHPLC.

Media components and supplements were purchased from Sigma-Aldrich, St. Louis, MO, unless otherwise indicated in the supporting information. A table of supplement and stock solutions with screening concentrations is also included in the supporting information. During modular optimization, all media were prepared in high throughput using an Opentrons OT-2 liquid handler (Opentrons, Brooklyn, NY, software version ≥3.16.1) using Openblend. Modular media blending code and instructions for setup and operation are provided in the Openblend package (https://github.com/abiedermann/openblend_public). For consistency, media used in final head-to-head comparisons were prepared in bulk and filter sterilized through a 0.2 *μ*m benchtop filter.

### Analytical procedures

Biomass was measured by optical density at 600 nm as described previously (Matthews et al., 2017a). An Agilent Bravo liquid handler was used to dilute samples prior to measurements of OD into the Tecan Infinite M200 Pro plate reader.

Reverse phase ultra-high performance liquid chromatography (UHPLC) analysis was performed on Agilent 1290 Infinity II UHPLC system controlled using OpenLab CDS software (Agilent Technologies, Santa Clara, CA). The concentration of protein was determined using a Poroshell 120 SB-Aq column (2.1 x 50 mm, 1.9μm) operated at 1.0 mL/min and 70°C (Agilent Technologies, Santa Clara, CA). Buffer A was 0.1% (v/v) TFA in water and buffer B was 0.1% (v/v) TFA, 0.5% (v/v) water in ACN. A gradient was performed as follows: 30% B for 1 min., 30-40% B over 3 min., 40-90% B over 0.5 min., 90% B for 0.5 min., 90-30%B over 0.5 min., and 30% for 1 min.; total method run time was 6.5 minutes. Sample injection volumes were 50μL. A diode array detector was set for absorbance detection at 214nm. Data analysis was completed using OpenLab CDS Data Analysis (Agilent Technologies, Santa Clara, CA).

Statistical analysis and DOE design was conducted using JMP (SAS Institute, Cary NC). Quadratic models were fitted using effect screening and non-significant terms (adjusted p-value > 0.01) were eliminated sequentially in order of decreasing adjusted p-value to avoid overfitting. Data was plotted using Prism 8.4.0 (GraphPad Software, San Diego, CA).

## Results

### Design of approach for modular media blending

We sought to develop an approach capable of identifying important, beneficial modifications for media tailored to a given phenotype of interest. We reasoned that key requirements for such an approach would be that it is fast and automatable, with minimal dependence on complex analytical assay development. Such features would enable routine application to any measurable phenotype of interest. In general, media blending allows both speed and low analytical complexity. We aimed to retain these features while minimizing the labor and constraints on compositions imposed by linear combinations of fully formed and unique media. We reasoned that diverse and flexible blends of media could be created by defining simple concentrated stock solutions as basic modules to combine further. These modules would comprise individual components or common subsets of components with compatible solubilities (e.g. YNB). If media components could be formulated in concentrated stock solutions that could be stored stably over time, then the components could be routinely and interchangeably combined and diluted to the desired final concentrations. This approach would yield a broadly applicable modular strategy for media blending amenable to conventional liquid handling automation.

To test the feasibility of this approach, we first assessed whether many common media components could be formulated in concentrated stable aqueous stock solutions. Using the CHO medium eRDF as a reference, we estimated the solubility of each component of this medium, using data from AqSolDB as well as other online sources (Combs, 2012; FSA Panel on Additives and Products or Substances used in Animal Feed (FEEDAP), 2011; Ritacco et al., 2018; Schnellbaecher, Binder, Bellmaine, & Zimmer, 2019; Sorkun, Khetan, & Er, 2019; Yamamoto & Ishihara, n.d.). We compared the estimated solubility of each media component to its concentration in eRDF and found that, individually, most media components are soluble at levels >10x higher than their eRDF concentration (**Figure 1A**). The existence of a wide range of commercially available concentrated supplements further supports this result: >50x concentrated solutions of amino acids, vitamins, lipids, and trace metal supplements are common and commercially available.

**Figure 1.**
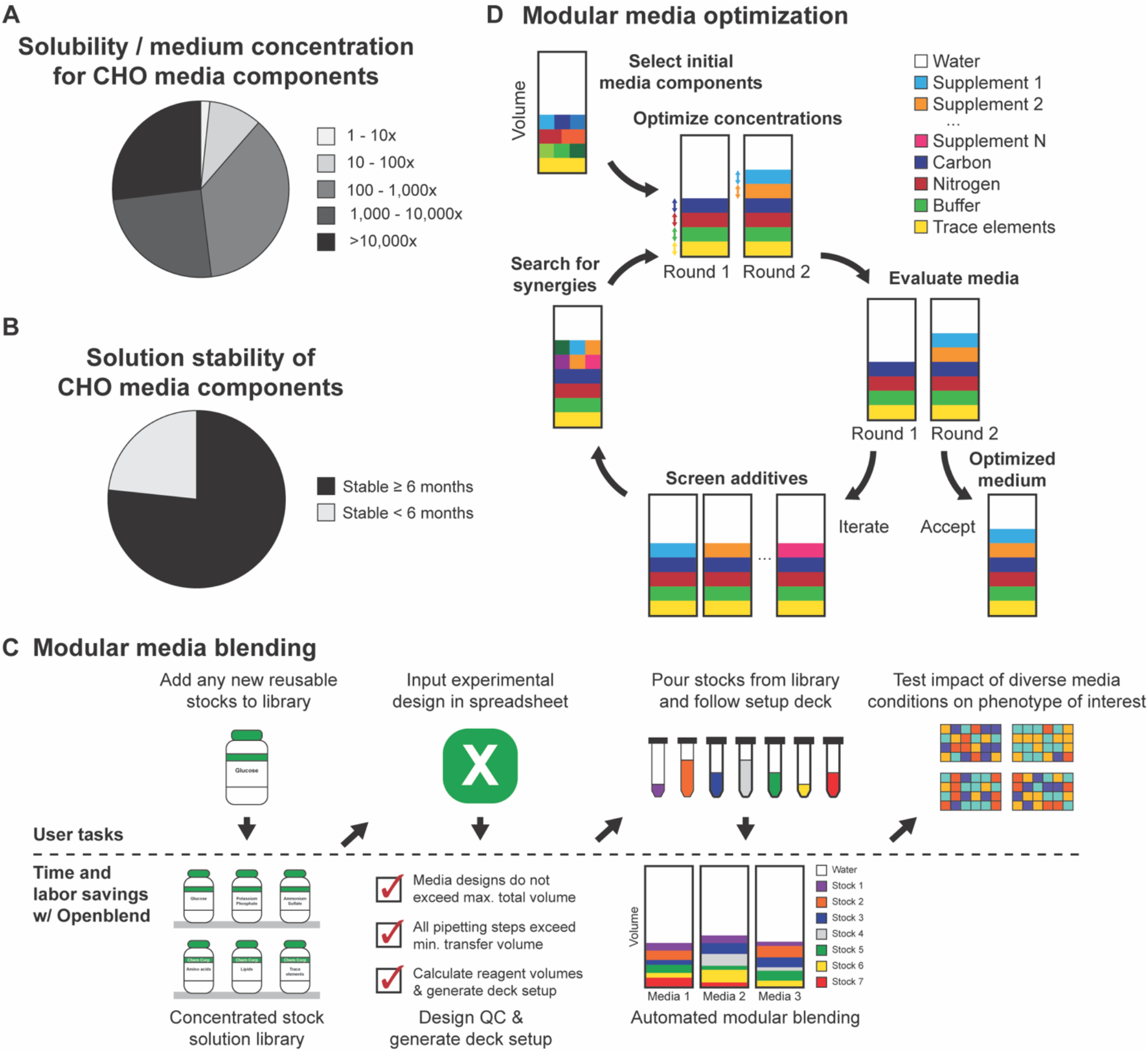
Modular media development can be broadly applicable, easily applied, and systematically executed to improve measurable phenotypes of interest. A) Estimate of the ratios of component solubility to their concentrations in medium demonstrates that most components are soluble at >10x their concentration in the CHO medium, eRDF. B) With the exception of some classes of medium components, such as vitamins, most media components can be formulated into solutions that remain stable for >6 months under proper storage conditions. C) Overview of time, labor and planning saved by using Openblend to automate modular media construction. D) Overview of a modular media optimization approach, which can be used to build an optimized medium for any measurable phenotype of interest systematically.

Next, we used the product information of commercially available supplements, literature sources, and inspection to estimate the percentage of eRDF media components that could be stored in stable solutions for >6 months. We estimated that over 75% of eRDF components met this criterion (**Figure 1B**). To address stability challenges caused by less stable components, we reasoned that less stable components or supplements, such as vitamins, could be prepared, aliquoted, and stored frozen for long-term storage (Schnellbaecher et al., 2019); these aliquots could then be thawed and used within a defined period to mitigate component instability and enable their integration into our modular blending strategy. Together, these solubility and stability data suggested that a modular approach to media development could be defined in this way to accommodate a range of new formulations easily.

We next automated the process for constructing media, using the Opentrons OT-2. We chose this liquid handler due to its low cost, reliability, and compatibility with simple formats for data input, such as Excel spreadsheets. We then created an open-source Python package, named Openblend, which simplified the media construction process by handling routine experimental design and execution steps (Figure 1C). Openblend creates an experimental design spreadsheet, specifying the number of 24 well plates, the desired media composition of each well, and stock solution names and concentrations. The script then checks the feasibility of the experimental design, ensuring that the total volume of each well will not exceed the target volume and avoiding the addition of sub-microliter stock solution volumes. If the design passes this assessment, the script then outputs a new spreadsheet containing the setup for the OT-2 deck and required volumes of stock solutions, providing a user with instructions on how to setup the OT-2 liquid handler. We found that our typical time to execute this script, setup the OT-2 and initiate plate building was ~15 minutes, and the time for the automated steps was about two hours.

Finally, we defined a modular approach for optimization to effectively leverage the Openblend tool (**Figure 1D**). Beginning from an initial basal medium, improved media are constructed through successive rounds of optimization. In each round, a library of media components and supplements are screened to identify beneficial additives. These additives are then screened in combination and over a range of concentrations to further optimize the performance of the medium. Each modular addition and optimization of additives can be guided simply by measurements of the phenotype of interest (e.g. biomass accumulation). This greedy approach to multi-dimensional optimization could continue iteratively until the resulting media met desired specifications, all available media components were explored, or no additional gains in performance realized.

### Application to Developing a Medium for Biomass Accumulation

To assess the utility of this blending-based approach, we next aimed to identify and optimize the concentration of media components beneficial for rapid biomass accumulation of *P. pastoris* in batch cultivation. We previously described a rich defined medium (RDM) (Matthews et al., 2017a), capable of high growth rates during biomass accumulation. One challenge encountered with this formulation, however, was that precipitates can form at higher pH values that require filtering during bulk preparations. Nonetheless, this medium provided a relevant comparison for assessing the medium realized with our new approach due to its prior demonstrated benefits relative to complex media. Following our modular approach, we improved biomass accumulation by optimizing the accumulated optical density at 600 nm after 24 hours of cultivation.

Algorithms for optimizing systems based on multiple dimensions are often sensitive to initial conditions used (Zakharova & Minashina, 2015). Given this potential confounding effect here, we tested first the effects of the types of carbon source, nitrogen source, and pH set point on biomass accumulation, using 1x YNB without amino acids or ammonium sulfate (YNB) to satisfy minimum requirements for the concentrations of trace elements. We conducted a full-factorial DOE using glycerol, glucose, and fructose as carbon sources; urea and ammonium sulfate as nitrogen sources; and potassium phosphate as a buffer with pH values of 5, 5.75, and 6.5. We selected initial concentrations of 40 g/L, 4 g/L urea or the N-mol equivalent for ammonium sulfate, and 10 g/L potassium phosphate, similar to values used in other media for *Pichia pastoris* (Matthews et al., 2017a). A least squares regression model, including individual, combination, and quadratic effects was fit to the log of optical density after 24 hours, a proxy variable for the average growth rate (R^2^ = 0.81). We determined that the two most significant model terms were the type of carbon source and the interaction of the nitrogen source with pH (**Figure 2A**). We found that cells grew significantly faster on metabolically related sugars (glucose and fructose) than on the polyol (glycerol) commonly used for Pichia during biomass accumulation (**Figure 2B**). This result affirms prior reports where glucose has been used for biomass accumulation of Pichia (Guo et al., 2012; Moser et al., 2017).

**Figure 2.**
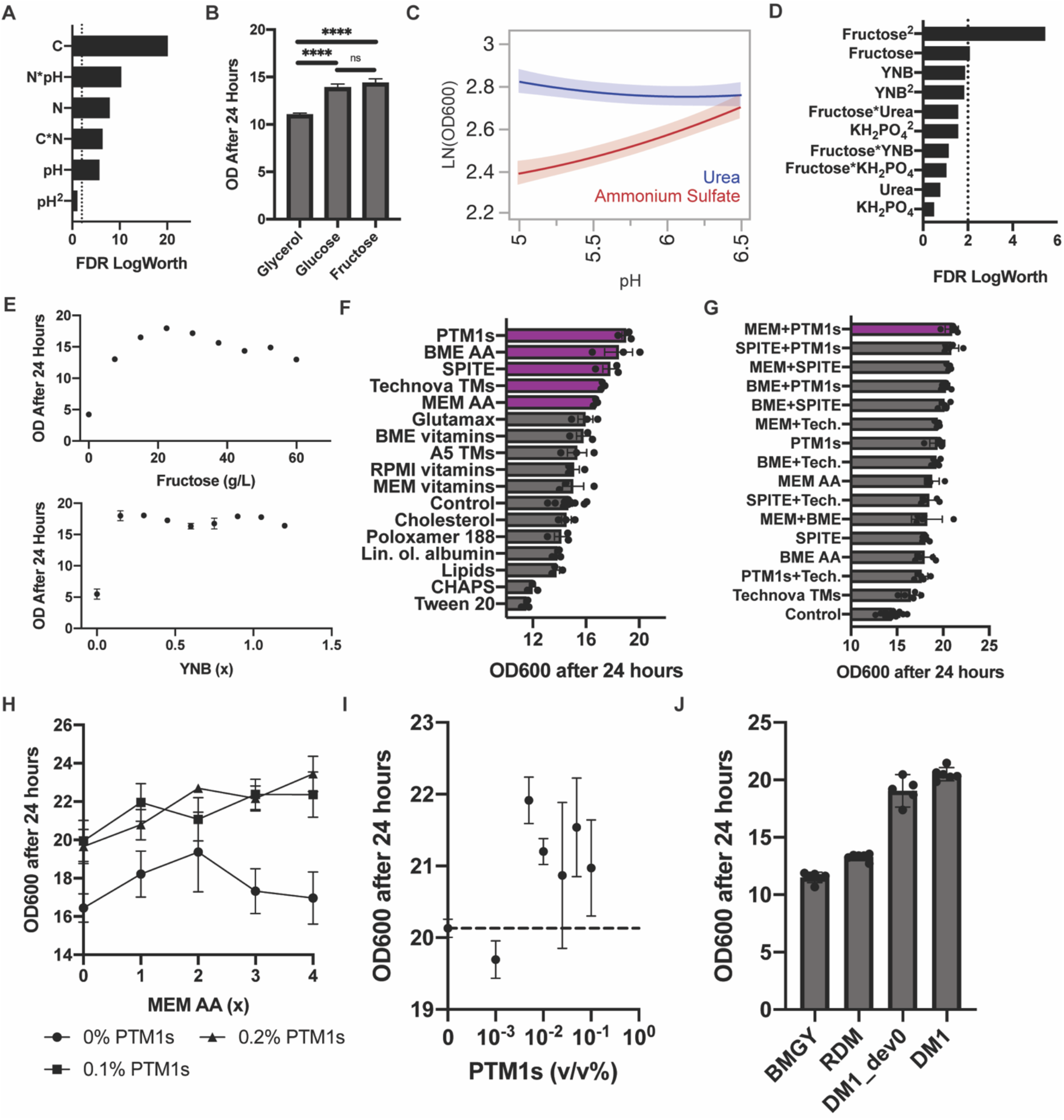
Modular development of a new biomass accumulation media for *P. pastoris*. A) Significance of carbon (fructose, glucose, glycerol), nitrogen (urea and ammonium sulfate), and pH choice (5, 5.75, 6.5) in a least square regression model fitted to a full factorial DOE. B) Fructose and glucose were found to result in significantly higher biomass accumulation after 24 hours of outgrowth than glycerol. C) Ammonium sulfate was found to be more pH sensitive than urea, as shown by the JMP sensitivity profiles during fructose feeding. D) Significance of terms in a least square regression model fitted to a full factorial DOE over fructose, urea, potassium phosphate, and YNB concentrations. E) 1-FAAT optimization of fructose and YNB concentration finds optimal outgrowth performance at a fructose concentration of 22.5 g/L and relative insensitivity over a wide range of YNB concentrations (0.15 to 1.2x). F) A media supplementation screen identified 5 beneficial supplements, related to trace element and amino acid supplementation. G) Further screening of beneficial supplement combinations identified synergistic amino acid and trace metal supplementation strategies. H) Comparison of the effect of MEM amino acid concentration on biomass accumulation at different PTM1 salts concentrations. I) Effect of the concentration of PTM1 salts on biomass accumulation in DM1_dev0 medium supplemented with 1x MEM AA. J) Head-to-head comparison of 4 v/v% glycerol BMGY, 4 v/v% glycerol rich defined medium, the initial defined biomass accumulation media (DM1_dev0), and the final biomass accumulation medium obtained after a full optimization cycle (DM1), demonstrates that DM1 leads to superior biomass accumulation.

The model also suggested that poor biomass accumulation during cultivation resulted from a combination of ammonium sulfate as a source of nitrogen with low buffer pH (**Figure 2B**). This outcome may result from the production of acidic species associated with cellular ammonium metabolism in the batch cultivation (Villadsen, 2015). Interestingly, the model indicated slightly greater biomass was achieved with urea instead of ammonium sulfate. The biomass accumulation of cultures grown with urea as a source of nitrogen were less sensitive to reduced pH values (~5). We observed, however, that cultivations at pH 5 showed extensive flocculation compared to those at 6.5. Given the insensitivity of urea-fed cultivations to buffer pH and the high solubility and potential for low-cost sourcing of fructose, we therefore chose to include fructose, urea, and a potassium phosphate buffer with a pH of 6.5 in our initial media formulation.

With this basal formulation determined, we next screened for concentration-dependent interactions of other key additives to the media and then optimized concentration-dependent parameters. Following the same approach for screening effects, we conducted a full factorial DOE over a broad range of media component concentrations: YNB (0.5, 1, 2x), fructose (10, 30, 50 g/L), urea (1, 4, 7 g/L), and potassium phosphate adjusted to a pH of 6.5 (4, 10, 16 g/L). The resulting model identified fructose as a concentration-sensitive parameter (R^2^=0.73) (**Figure 2D**). Terms involving the concentration of YNB were also highly ranked, but not statistically significant. No significant interactions between components were identified in the model. We therefore sought to better understand the concentration dependence of fructose and YNB independently (**Figure 2E**), over an 8-fold range of concentrations. As expected, biomass accumulation was highly sensitive to fructose concentration, with an optimum around 22.5 g/L of fructose. The concentration of YNB had minimal effect on biomass accumulation; the presence of trace elements supplied by YNB, however, was essential to growth. Based on these results, we chose concentrations of 22.5 g/L fructose, 1x YNB, 7 g/L urea, and 10 g/L potassium phosphate buffer. We reasoned that although biomass accumulation was relatively insensitive to the concentrations of YNB and urea, higher concentrations could provide improved media depth in future applications. We named this basal formulation DM1_dev0.

We next assessed what additional media components could improve biomass accumulation. To test over 60 different components individually would require over 60 individual solutions. Such an approach would scale linearly with new components; instead, we chose to screen groups of related components, using commercially available pre-mixed supplements. We compiled a library of 16 commercial supplements and industrially-relevant surfactants containing more than 60 unique components and screened their individual effect on biomass accumulation after 24 hours. In this way, we reasoned we could efficiently identify critical classes of components related to the phenotype of interest and potentially deconvolve specific individual additives of interest by inference. We used the recommended concentrations of each supplement as supplied in product information, or critical micelle concentrations, and prior knowledge for broad classes in yeast media to set reasonable screening concentrations (Supporting Information). We identified five beneficial and two detrimental supplements that significantly impacted biomass accumulation (p_adj_ < 0.02; 1-way-ANOVA) (**Figure 2F**). In general, the results suggest that supplementation with amino acids and trace metals were beneficial for accumulating biomass, while two surfactants, Tween 20 and CHAPS, were detrimental. For this phenotype, the effects of vitamin and lipid supplements were minor; supplements from either supplement category were not significantly beneficial or detrimental to biomass accumulation. Our earlier experiments suggest that vitamins are essential but concentration agnostic (**Figure 1E**), while lipid supplementation provides no clear benefit for biomass accumulation.

Based on these results, we chose to test whether combinations of supplements of amino acids and trace salts could yield synergistic improvements in biomass accumulation. We screened pairwise combinations of the five beneficial supplements of mixed composition and ranked the performance of our supplementation strategies (**Figure 2G**). A combination of 1x MEM amino acids with 0.1 v/v% PTM1 salts resulted in the highest yield of biomass, though we observed strong performance from other combinations of amino acid and trace metal supplements. Based on these data, we chose to add MEM amino acids and PTM1 salts in our basal medium and optimized their concentrations (**Figure 2H**).

Based on these results, we elected 0.1 v/v% PTM1 salts and 1x MEM amino acids, in order to balance the moderate benefits and potentially high costs of amino acids. We found, however, that the inclusion of the PTM1 salts in liter-scale preparations produced fine precipitates, which can impede sterile transfers in use. To overcome this challenge, we screened a broad range of PTM1 salts concentrations to identify the minimum concentration required for improved outgrowth performance (**Figure 2I**). We found that PTM1 addition at concentrations as low as 0.0005 v/v% led to increased biomass accumulation. We therefore revised our PTM1 salts concentration to 0.01 v/v%, a concentration high enough to obtain the benefits of PTM1 supplementation without inducing precipitate formation. This formulation we named DM1.

Completing this series of optimizations with our iterative modular approach to define a new formulation of medium, we then compared with other common media used to grow *P. pastoris*. We evaluated the performance of this new optimized medium (DM1) relative to the unsupplemented basal medium (DM1_dev0), the rich defined medium (RDM) we had previously developed, and a common medium 4 v/v% glycerol BMGY. We found that DM1 yielded the highest biomass accumulation, with significantly higher biomass accumulation relative to RDM and BMGY (**Figure 2J**). This result demonstrates the utility of our modular strategy here for media development that yielded an improved formulation for biomass accumulation compared to other common media with minimal time and labor investment, and without requiring complex analytical methods like mass spectrometry or RNA-sequencing.

### Identifying media conditions important to heterologous protein production in *K. phaffii*

In addition to the time and labor savings of modular media development, our proof-of-concept experiments demonstrated that this approach creates a flexible medium that can be rapidly adapted to new growth phenotypes, as well as a data package that the identifies media conditions important to the phenotype of interest. We reasoned that these additional benefits could be particularly relevant for optimizing production of heterologous proteins. Understanding which media components contribute most significantly to productivity could improve culture performance and help identify important metabolic pathways or physiological effects for further study.

To develop a medium for improved production of a recombinant protein, we chose to use a strain engineered to secrete a rotavirus-derived subunit vaccine component, VP4-P[8], as a model protein. We have previously demonstrated that this viral antigen can be expressed at high titer under the control of the methanol-inducible pAOX1 promoter in BMMY media (Dalvie et al., 2020). Similar to our initial approach to optimize a medium for growing biomass, we first determined and optimized the concentrations of the sources for carbon and nitrogen, along with the pH. The expression of P[8] in the strain tested uses the methanol-dependent pAOX1 promoter for inducible expression, so we selected methanol as the initial carbon source. We then examined the impact of the source of nitrogen and buffer pH on titer. We conducted a full-factorial DOE using identical concentrations as those used to create a medium for accumulating biomass. The resulting model was visualized by ranking combinations of sources of nitrogen and buffer (**Figure 3A**). The effects showed no interaction between these two factors. Urea was again identified as the preferred source of nitrogen while higher pH values led to improved secreted P[8] productivity. Unlike biomass accumulation, this pH dependence was observed across both nitrogen sources.

**Figure 3.**
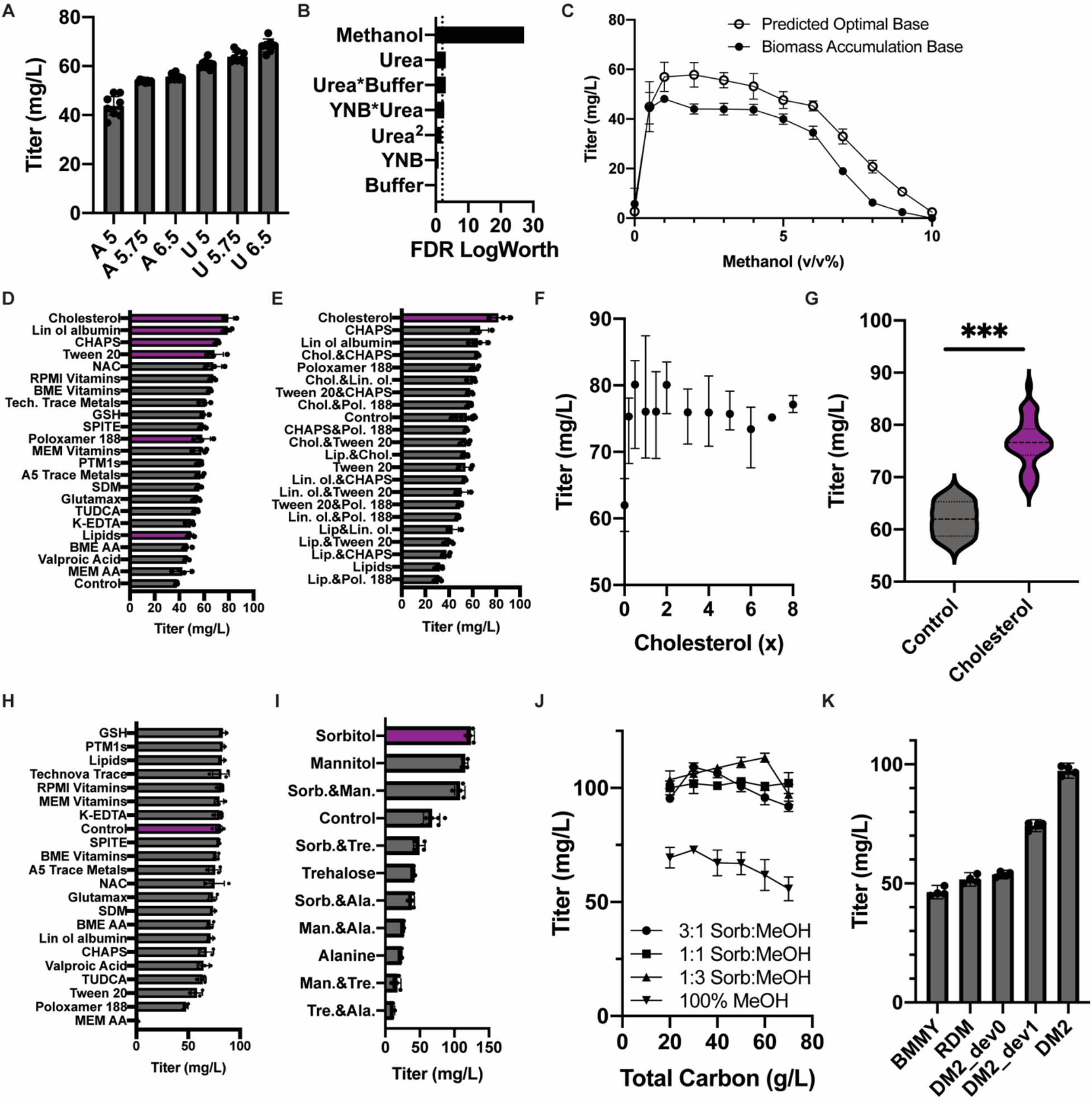
Modular development of a media for heterologous protein production in *P. pastoris*. A) Initial full-factorial screen of nitrogen source choice and buffer pH demonstrates that urea is preferred over ammonium sulfates and high buffer pH is preferred over lower values. B) A full-factorial concentration optimization identified methanol as the most concentration dependent variable. Other components in the base media were predicted to affect productivity with much lower levels of significance. C) Evaluation of the effect of methanol concentration on P[8] titer, using two different base media (urea, buffer, and YNB concentrations): the biomass accumulation base medium and the optimal base media composition predicted by our concentration DOE. D) Ranking of supplements according to their effect on P[8] titer. Supplements related to membrane fluidity or lipid metabolism ranked highly. E) Evaluation of combinations of lipid and surfactant supplements confirmed that cholesterol supplementation leads to the greatest improvement in P[8] titer. F) Concentration optimization of cholesterol demonstrated low concentration dependence, with similar performance observed over a 40-fold range (0.2-8x). G) Comparing cholesterol-free and cholesterol-supplemented cultures fed at various concentrations demonstrates that cholesterol supplementation results in a significant ~25% improvement in P[8] titers (p<0.001). H) No significantly beneficial supplements were observed when repeating the supplementation screen. I) Screening supplementation of 20 g/L of co-fed substrates individually or in 1:1 combinations by mass identified sorbitol supplementation as highly beneficial to P[8] titer. J) Examination of the effect of co-feed ratio and total carbon concentration on titer in DM2_dev1 supplemented media. K) Comparison of P[8] titer obtained with DM2 to previous iterations and other common *P. pastoris* media demonstrates a ~2x improvement in P[8] titer, relative to 1 v/v% methanol RDM and 1 v/v% methanol BMMY.

We next applied the same DOE to identify important concentration-dependent interactions that impact the production of P[8]. Unsurprisingly, the concentration of methanol was the most important factor, with possible minor effects from other components (**Figure 3B**). We decided to screen further a 20-fold range in methanol concentrations using two formulations for remaining media components—the one determined for optimal cell growth (DM1) and the optimal base media formulation predicted by the quadratic model here (2x YNB, 1 g/L urea, 4 g/L potassium phosphate adjusted to a pH of 6.5). We found that production was relatively insensitive for concentrations of methanol ranging from 1-4 v/v%, with an optimum around 2% (**Figure 3C**). We postulated that the rapid decline in productivity observed in these milliliter-scale cultures using concentrations >6 v/v% methanol was likely due to excess formation of toxic metabolic byproducts such as formaldehyde and hydrogen peroxide (Wakayama et al., 2016). Interestingly, the predicted optimal medium from this set of studies outperformed the medium we determined for accumulating biomass, suggesting that certain components of the basal medium may benefit protein expression more than cellular growth and underscores the value of optimizing media for specific phenotypes of interest. Based on these data in total, we defined a basal medium for production including 2x YNB, 2 v/v% methanol, 1 g/L urea, and 4 g/L potassium phosphate buffer adjusted to a pH of 6.5 (DM2_dev0).

Next, we examined which supplements could improve the performance of DM2_dev0. We added three chemical chaperones (TUDCA, sodium deoxycholate monohydrate (SDM), and valproic acid) (Kuryatov, Mukherjee, & Lindstrom, 2013; Uppala, Gani, & Ramaiah, 2017), two antioxidants (reduced glutathione (GSH) and N-acetyl cysteine (NAC)), and the chelator, K-ETDA, to the list of 16 supplements included in our original screen defined for biomass accumulation. Concentrations for these components were chosen based on product specifications, literature data, and prior experience (Supporting Information). Many of the 22 supplements screened improved production of P[8] (**Figure 3D**). The top four ranking supplements comprised surfactants or lipids, which could modulate membrane fluidity and lipid metabolism (Butler, Huzel, Barnab, Gray, & Bajno, 1999; Degreif, Cucu, Budin, Thiel, & Bertl, 2019; Ritacco, Frank V; Yongqi Wu, 2018).

We then screened combinations of lipid supplements and surfactants to identify potential synergistic effects. We ranked the individual supplements and their combinations (**Figure 3E**) according to the measured titers of P[8]. We found that the addition of a cholesterol-rich supplement yielded the highest secreted titers of P[8] (~50% improvement compared with supplement-free condition in initial screens). Interestingly, a synthetic cholesterol supplement alone did not substantially improve performance, suggesting the benefit results from a combination of fatty acids and surfactant components in the supplement (**Supporting Information**). This conclusion is consistent with similar improvements observed from other supplements, such as linoleic acid-oleic acid-albumin (**Figure 3D**).

Since no other synergistic effects were observed in the combination screen, we assessed the dependence of titer on the concentration of the cholesterol-containing supplement identified (**Figure 3F**). Similar to our observations with cellular YNB used in the outgrowth media, we found that concentrations of the supplement as low as 0.2 v/v% were beneficial for protein expression, but that production was relatively insensitive to concentration (**Figures 3F, 3G**). We then directly compared the supplemented medium to the original composition; the new supplemented media provided a 25% improvement in titer (p = 0.0006, one-tailed Welch’s T test). This new formulation with 1x cholesterol supplement, which we named DM2_dev1, was the result of one cycle of optimization using our method.

Components of the cholesterol supplement included fatty acids, cholesterol, and cyclodextrin, which are all are known to modulate membrane fluidity, a key parameter in vesicle trafficking (Cooper, 1978; Degreif et al., 2019; Mahammad & Parmryd, 2015). We reasoned that the addition of this supplement could therefore have synergistic effects with other supplements, but did not find any further supplementation that improved P[8] titers within our original screen (**Figure 3H**). We, therefore, considered if there could be additional classes of beneficial supplements, absent from the original screen. Previous experiments demonstrated that P[8] productivity is highly sensitive to methanol concentration (**Figure 3C**), so we wondered whether further modulation of central carbon metabolism could yield additional productivity gains.

Modification of central carbon metabolism is best accomplished by feeding cells alternative carbon sources, either entirely or as co-feeding substrates. Four co-fed substrates have previously been shown to be non-repressive of pAOX1: sorbitol, mannitol, trehalose, and alanine (Inan & Meagher, 2001). These substrates can be co-utilized with methanol without repressing the pAOX1 promoter, which controls expression of P[8]. We hypothesized that the introduction of supplemental carbon sources could enable further optimization of central carbon metabolism. We screened co-fed substrates individually and in 1:1 combinations at a total concentration of 20 g/L (a concentration similar to the optimal fructose and methanol concentrations observed in previous carbon source optimizations) (**Figure 2E,3C**). Sorbitol co-feeding had the most beneficial effect, resulting in a ~80% increase in P[8] titer (**Figure 3I**). Mannitol supplementation was also beneficial (~70% increase), while alanine and trehalose co-feeding were detrimental to productivity. While co-feeding carbon sources led to increased biomass yield during production, these differences did not account for the improved titer, as improvements in specific productivity (q_p_) of ~60% and ~45% were also observed for the sorbitol and mannitol co-fed conditions, respectively (**Supporting Information**). Based on these data, we chose to include sorbitol as a supplemental carbon source for further study.

The addition of a supplemental carbon source could significantly impact central carbon metabolism. We, therefore, wondered how the inclusion of sorbitol might impact the optimal carbon feeding strategy. Examining total carbon source concentrations from 20 – 70 g/L, we compared the performance of cultures co-fed with sorbitol:methanol ratios of 3:1, 1:1, and 1:3 to a methanol-only control (**Figure 3J**). All co-fed conditions outperformed the methanol-only control, suggesting that the presence of sorbitol is highly beneficial for producing P[8]. The titer was relatively insensitive to sorbitol:methanol ratios and carbon concentrations. Based on the data, we decided to use 2 v/v% methanol and 20 g/L of sorbitol for the final sorbitol-supplemented media named DM2.

Finally, we compared the P[8] titer obtained using DM2_dev0, DM2_dev1, and DM2 to other common production media for *P. pastoris*: BMMY and RDM. We found that DM2 led to a ~2x improvement in P[8] titers, relative to BMMY and RDM, up to 97 ± 2 mg/L.

## Discussion

Here we have implemented a novel and broadly applicable approach for media development that relies on rapid, automated construction of diverse media from defined modules of components. We demonstrated the utility of this approach by developing two new media for two phenotypes of interest in the heterologous production of proteins by yeast, namely biomass accumulation and secreted production. We systematically identified and optimized the concentration of media components important to each phenotype of interest. Importantly, defining these new formulations of media did not require advanced analytical capabilities and required minimal experimental time to assess more than 360 total formulations during two to three rounds of optimization for each.

Our optimized formulations affirmed the importance of lipid-related components for maximizing titers in *Pichia pastoris* cultivations. The importance of optimizing membrane fluidity or lipid metabolism has been well established in CHO and appears to be key to optimizing heterologous protein secretion in *P. pastoris* cultivation as well (Clincke et al., n.d.; Ritacco et al., 2018; Zhang, Wang, & Liu, 2013).

Modular media blending has four advantages over existing methods. First, the use of common stock solutions and supplements to formulate media reduces initial labor required for new experiments or optimizations ~15 minutes per experiment, making parallel testing of multiple hypotheses efficient and requires less resources overall. Here, we created 30 stock solutions, and evaluated >360 unique media compositions, without manual preparation of individual media or extensive blending calculations or planning. Most of these solutions could be readily reused in future experiments to optimize for new phenotypes of interest. Second, our method requires minimal knowledge of the host organism *a priori* and could, in principle, be applied to any measurable phenotype of interest. We anticipate that this method could be used to optimize other phenotypes of interest, such as glycosylation profiles. Third, our method provides certain practical advantages, including minimal requirements for analytical characterization and rapid identification of component interactions that lead to solubility challenges. These traits make it possible to learn about formulations that may lead to extensive precipitates like those encountered with our rich defined medium formulation (**Figure 4A**). Finally, modularly constructed media, such as DM2, can be ~70% pure water with low osmolarity, leaving volumetric and osmotic space for future modifications to accommodate new or related phenotypes of interest (**Figure 4B**).

**Figure 4.**
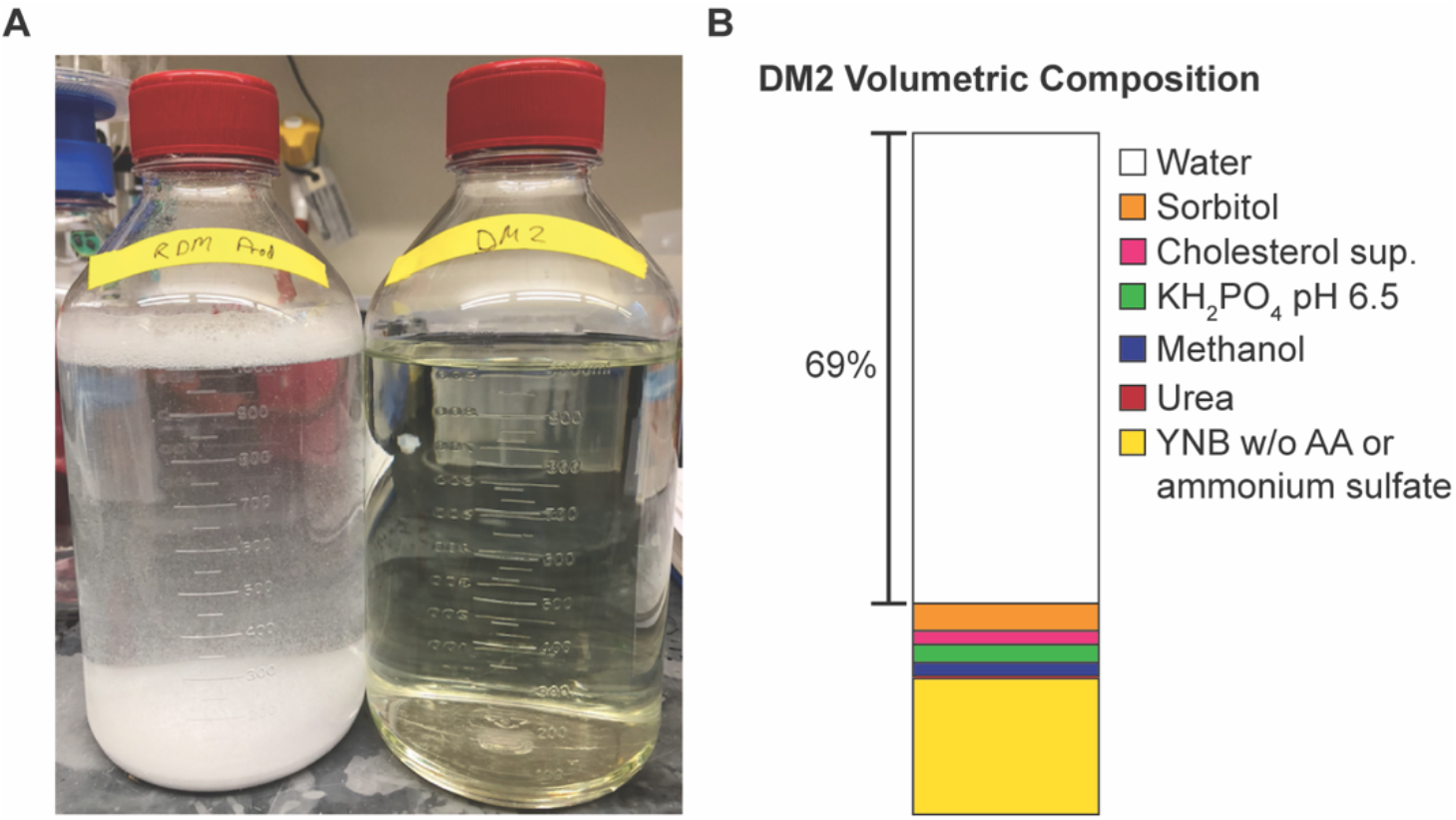
Comparison of DM2 to rich define medium. A) Comparison of precipitate formation during construction of RDM (left) and DM2 (right) media. Adjusting the pH of RDM to 6.5 results in significant formation of white precipitate. No precipitate formation is observed in DM2. B) Relative volumes of stock solutions and pure water needed to construct DM2. Pure water addition accounts for 69% of DM2 volume, demonstrating that there is substantial room for further supplement exploration and development. When separated into simple stock solutions, DM2 can be 3x concentrated.

We also acknowledge certain limitations in the present study that may be addressed in future work. First, while modular media development identifies components key to the optimization of the phenotype of interest, additional media optimization effort may be necessary to translate these learning in batch cultivations to scaled-up fed-batch or perfusion operation, where additional variables such as supplemental feed composition and feeding schedule must also be considered. In principle, modular media construction could be applied to high-throughput scale-down cultivation models, such as Ambr250s. Second, our approach for optimization relies on greedy algorithms tailored to create a new media for a single phenotype of interest; however, given the vast explorable media space it is possible to find a local optimum. Further metabolic or -omic modeling techniques could be employed to guide broader exploration of media space, co-optimize multiple phenotypes, or facilitate biologically informed optimization, albeit with more complex experimental and computational requirements (Matthews, Kuo, Love, & Love, 2017b; Mohmad-Saberi et al., 2013). Third, our current method used commercially available supplements, but in practice, beneficial supplements could be simplified by using individual components, to facilitate more biological inferences and aid development of improved host-specific supplements. Finally, initial screens to identify beneficial supplements rely on reasonable choices of initial concentrations for screening. These currently require prior knowledge from the literature or commercial sources; with further use in the community of the Openblend approach, it is possible additional sharing of knowledge could help inform further developments.

The improved speed and accessibility of in-depth media development experiments enabled by modular media construction could help improve expression of many classes of proteins in laboratories and discovery centers that have not traditionally had access to such capabilities. Since many lead candidates for new therapeutic proteins begin in small biotech firms and academic labs, early-stage improvements in productivity could help advance more proteins towards the clinic simply by facilitating access to larger quantities of proteins for initial research and non-clinical studies. In more established companies, the ability to make rapid improvements to existing media may enable faster product development timelines and could reduce manufacturing costs overall. Rapid identification and optimization of sensitive media components could also enable easier adoption of a range of industrially relevant alternative hosts, resulting in further manufacturing flexibility and potentially cost savings (Coleman, 2020).

## Acknowledgements

The authors acknowledge Danielle Camp for program coordination. This work was funded by the Bill & Melinda Gates Foundation (Investment ID INV-002740). The content is solely the responsibility of the authors and does not necessarily represent the official views of the Bill & Melinda Gates Foundation.

A.M.B., I.R.G., K.R.L., and J.C.L. conceived and planned experiments. A.M.B. conducted media development experiments. S.R.A. developed and maintained the RP-UPLC assay. A.M.B. performed analytical characterization. A.M.B., K.R.L., and J.C.L. wrote the manuscript. J.C.L. and K.R.L. designed the experimental strategy and supervised analysis. All authors reviewed the manuscript.

