## Supplemental Figures for "Modular development enables rapid design of media for alternative hosts" (from another version of this preprint: version 1, 2021-04-30)

**Code:** The Openblend package at: https://github.mit.edu/lovelab/openblend

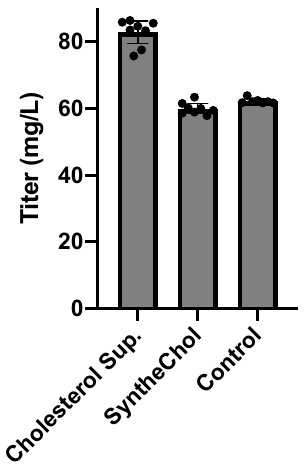

**S1**. Synthetic cholesterol supplementation alone does not elicit a productivity enhancement comparable to the multicomponent cholesterol supplement

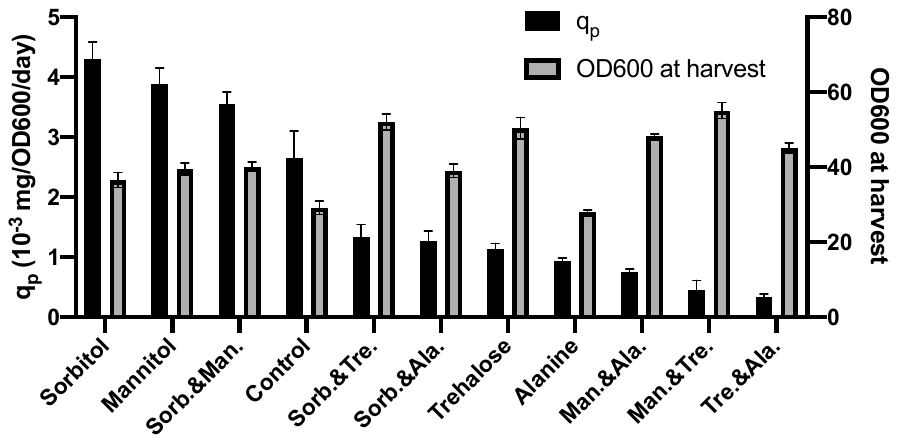

**S2**. Specific productivity and OD600 at harvest for co-feed supplement screen
